# Times to Key Events in the Course of Zika Infection and their Implications for Surveillance: A Systematic Review and Pooled Analysis

**DOI:** 10.1101/041913

**Authors:** Justin Lessler, Cassandra T. Ott, Andrea C. Carcelen, Jacob M. Konikoff, Joe Williamson, Qifang Bi, Nicholas G. Reich, Derek A. T. Cummings, Lauren M. Kucirka, Lelia H. Chaisson

**Author notes:** denotes equal contribution (Cassandra T. Ott, Andrea C. Carcelen, Jacob M. Konikoff).

## Abstract

**Background:** Evidence suggests that Zika virus has driven a 10-fold increase in babies born with microcephaly in Brazil, prompting the WHO to declare a Public Health Emergency of International Concern. However, little is known about the natural history of infection. These data are critical for implementing surveillance and control measures such as protecting the blood supply.

**Methods:** We conducted a systematic review and pooled analysis to estimate the distribution of times from Zika infection to symptom onset, seroconversion, and viral clearance, and analyzed their implications for surveillance and blood supply safety.

**Results:** Based on 25 cases, we estimate the median incubation period of Zika virus infection is 5.9 days (95% CI: 4.4-7.6), and that 95% of those who do develop symptoms will do so by 11.1 days post-infection (95% CI: 7.6-18.0). On average seroconversion occurs 9.0 days (95% CI, 7.0-11.6) after infection, and virus is detectable in blood for 9.9 days (95% CI: 6.8-21.4). In 5% of cases detectable virus persists for over 18.9 days (95% CI: 12.6-79.5). The baseline (no screening) risk of a Zika infected blood donation increases by approximately 1 in 10,000 for every 1 per 100,000 person-days increase in Zika incidence. Symptom based screening reduces this by 7% (RR 0.93, 95% CI 0.86-0.99), and antibody screening by 29% (RR 0.71, 95% CI: 0.28-0.88).

**Conclusions:** Symptom or antibody-based surveillance can do little to reduce the risk of Zika contaminated blood donations. High incidence areas may consider PCR testing to identify lots safe for use in pregnant women.

## INTRODUCTION

The explosion of Zika cases in Central and South America, combined with growing evidence that the virus is responsible for an epidemic of microcephaly in Brazil, has prompted the World Health Organization (WHO) to declare a Public Health Emergency of International Concern.^1^ As of February 29, 2016, there have been at least a half-million Zika virus infections in the Americas.^2,3^ Although clinical disease is generally mild or asymptomatic,^4^ there is increasing evidence of a link between Zika virus infection and severe microcephaly in infants born to women infected during pregnancy, including a 10-fold increase in microcephaly cases in Brazil in the wake of the 2015 Zika epidemic^5^. Zika virus infection has been linked to Guillain-Barre in adults.^5,6^

The severity of these complications highlights the need to protect pregnant women from infection and to ensure that blood supplies remain safe both in areas experiencing ongoing Zika virus transmission and in locations with travelers returning from affected areas. That large proportion of Zika infections that remain asymptomatic,^4^ inadequacy of current diagnostics, and uncertainties regarding the duration of viremia and viral shedding have raised concerns about the potential threat of transmission through blood transfusion. In a 2013-2014 outbreak in French Polynesia, researchers found that 3% of asymptomatic blood donors were infected with Zika virus,^7^ and several cases of possible Zika transmission through blood transfusion are currently being investigated in Brazil.^8^ As a result, some agencies now recommend halting blood donations in areas with active Zika transmission.^9,10^ If implemented, these bans could result in severe blood supply shortages. Research to determine the duration of viremia and time to antibody seroconversion is therefore vital to quantify the risk to blood supplies, and develop efficient strategies for protection. Furthermore, more detailed estimates of key distributions in the natural history of Zika virus infection, including the incubation period and probable infectious period, are essential to designing evidence based surveillance systems and informing public health policy.^11,12^

In order to better characterize the natural history of Zika infection and inform disease prevention, surveillance, and blood supply safety, we performed a systematic literature review and pooled analysis of available data to estimate the incubation period, time to seroconversion, and length of shedding of Zika virus in infected populations.

## METHODS

The systematic review was conducted according to the Meta-analysis of Observational Studies in Epidemiology (MOOSE) guidelines^13^ and the Preferred Reporting Items for Systematic Review and Meta-Analyses (PRISMA) guidelines^14^ where applicable (see Supplemental Material).

### Search strategy and selection criteria

We searched PubMed, Scopus, and Web of Science on February 8,2016 for all publications containing the word “Zika” in any field, with no restriction on date of publication or language. The search was updated on February 25, 2016 to identify additional relevant publications.

We included publications that provided information on time of exposure to Zika virus and 1) time of symptom onset, 2) time of sample collection for virologic Zika virus testing (e.g. PCR or culture) and test results (positive/negative), and/or 3) time of collection of samples for antibody testing and test results (positive/negative). We excluded publications if they did not provide sufficient information to determine a bounded time of exposure to Zika virus, contained no data from humans, were not in English or French, or reported only perinatal transmission of Zika virus.

Two reviewers (CTO, LHC, JW, AC) independently screened titles and abstracts for relevance. We excluded publications from full text review if they were not about Zika virus or if they definitively met one of the exclusion criteria. Two reviewers (CTO, LHC, JW, AC) independently performed full text reviews to identify publications with sufficient data for analysis; we contacted authors via email to obtain additional information for publications that were relevant but did not provide sufficient data for analysis. Discrepancies were resolved by discussion and consensus.

### Data abstraction

Two reviewers (CTO, LC, JW, AC) independently abstracted data using a standardized form and resolved discrepancies by discussion and consensus. We abstracted data necessary to estimate: 1) the incubation period of Zika virus, 2) the time and duration of viral shedding, and 3) the time to antibody seropositivity. We reviewed text, tables, and figures for information that allowed us to bound the time of: 1) exposure to Zika virus, 2) symptom onset, 3) collection of samples for Zika virus testing, and 4) collection of samples for antibody testing. For all virologic and serologic samples, the reported test result was recorded, and IgM specific serologies were noted when available. When possible, the exact timing of events was used, otherwise timing of the event was bounded based upon available information (e.g., travel dates to Zika endemic regions). We further recorded basic demographic characteristics, the type of sample collected (e.g., blood, urine), and, when available, the mode of transmission.

Extracted data was used to construct a data set bounding the time of key events. The time of Zika infection was bounded by the earliest and latest potential time of Zika virus exposure consistent with the case report. When no latest time of exposure could be determined (e.g., the case developed symptoms while in a Zika endemic area) the latest possible time of symptom onset was considered to be the latest possible time of exposure, the most conservative assumption we could make. Time of symptom onset was bounded based on the case report, and in most cases was specified to the nearest day. The earliest possible time of seroconversion was considered to be immediately after the last negative serological test, and the latest possible time of seroconversion was immediately before the first positive serological test. If there was only a positive serological test, the earliest possible time of seroconversion was considered to be the same as the earliest possible time of exposure; when only negative tests were performed, time to seroconversion was considered to be right censored. Similarly, the earliest possible time of viral clearance was the time of the last positive test (by PCR or viral culture), and the latest was the time of the first subsequent negative test. Missing virologic tests were treated the same as in the serological data.

### Statistical Methods - Pooled Analysis

Bounding periods were used to construct doubly interval censored data sets for each distribution,^15^ and distributions were fit using an adaptation of techniques previously described.^15,16^ Briefly, MCMC techniques were used to simultaneously fit the incubation period distribution (log-normal), distribution of time to IgM seroconversion (Weibull), and time to viral clearance (Weibull) to the doubly interval censored data. Given a time of infection the distributions were considered to be independent. The mean incubation period, and the times by which we expect 5%, 25%, 50%, 75%, and 95% of those who do develop Zika symptoms to become symptomatic were estimated. Full details are available in the supplemental appendix.

### Statistical Methods - Blood Supply Safety

The impact of key distributions on blood supply safety was calculated assuming a constant incidence rate. The number of possibly infected donors per 100,000 if no screening occurs was calculated as: (daily incidence rate per 100,000) × (mean time to viral clearance). This estimate was adjusted for symptom based screening based on mean time to symptom onset, assuming that 80% of the population remains asymptomatic. The effect of serological based screening was calculated based on the mean time to the first of seroconversion or viral clearance (assuming independence), as the former cases would be successfully screened, and the latter would no longer be infectious. We assumed that any screening protocol would treat equivocal IgM ELISA results as being seropositive. Full equations are provided in the supplement.

Sensitivity to key model assumptions was assessed (see Supplement for results).

Analyses were performed using JAGS and the R Statistical Language.^17,18^ All data and code is available on GitHub (https://github.com/HopkinsIDD/ZikaLitReview).

## RESULTS

*Systematic Review Results* We identified 964 articles discussing Zika indexed by Pubmed, Scopus and/or Web of Science as of February 25, 2016 (Figure 1). After abstract and title screening, 846 articles were excluded based on predetermined criteria (480 duplicates, dead links, or non-English or French; 366 not about Zika in humans or lacking appropriate data). Among 118 articles selected for full text review and possible data abstraction, 86 did not have sufficient exposure information or dates of onset to estimate key distributions, and 11 reported suspected perinatal transmission. Authors were contacted for 4 articles lacking sufficient information on one or more cases; additional information was returned on one case, but we were still unable to bound the time of exposure. We extracted data from 21 articles that provided information on 25 unique Zika cases (Table 1). The analytic data set included: 25 individuals with a bounded time of symptom onset, 49 virologic tests on 22 individuals, and 62 serologic tests on 22 individuals.

**Figure 1:**
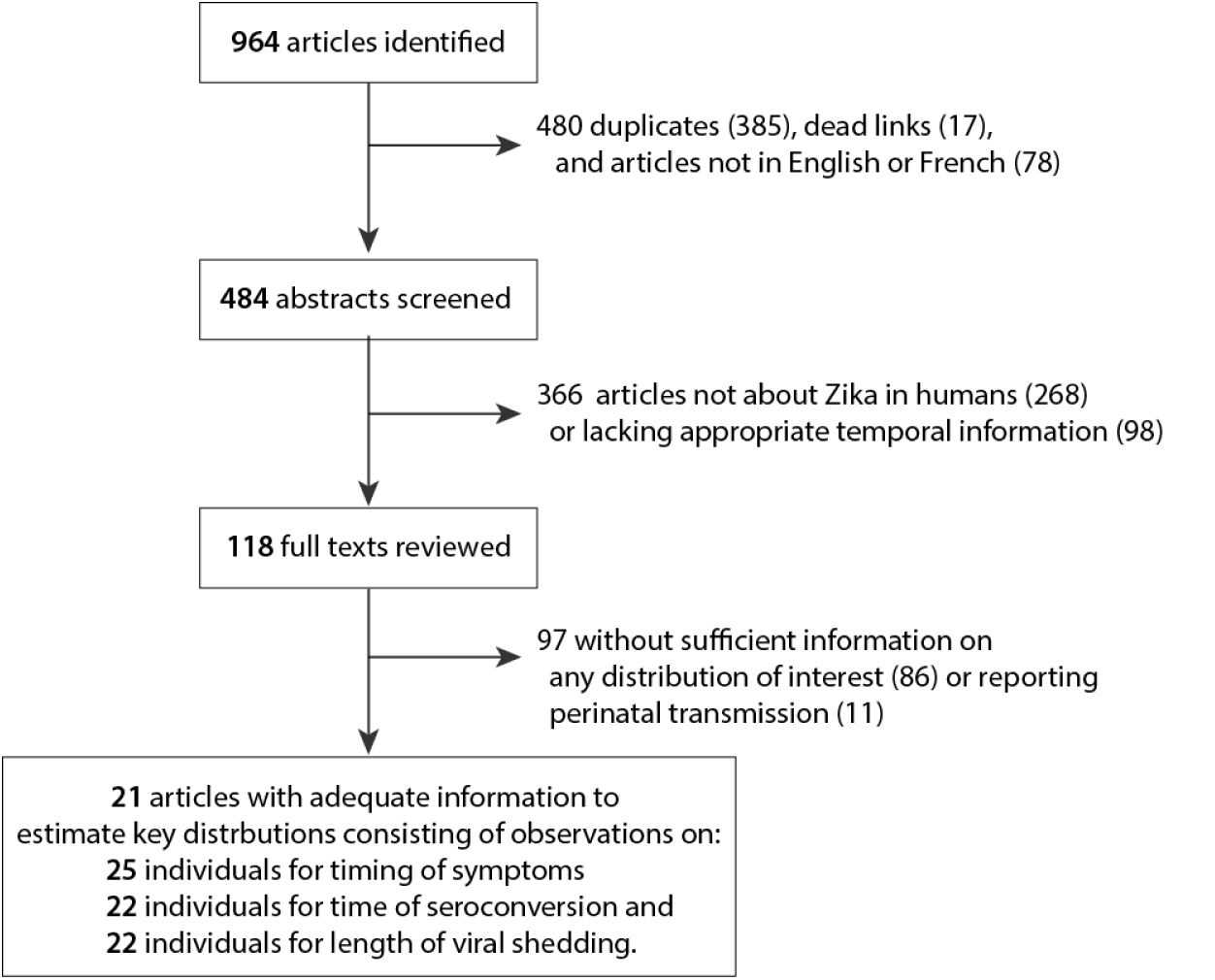
Systematic review process.

**Table 1.** Characteristics of Zika cases included in pooled analysis (N=25)

| First author (year) | Age | Sex | Place of origin | Probable location infected | Year exposed | Exposure window (days) | Days to symptom onset (min-max) | Days to seroconversion (min-max) | Days to viral clearance<sup>a</sup> (min-max) |
| --- | --- | --- | --- | --- | --- | --- | --- | --- | --- |
| Bearcroft (1956)<sup>19</sup> | 34 | Male | Europe | Nigeria<sup>b</sup> | – | <1 | 3-4 | 4-9 | >6<sup>c</sup> |
| Chen (2016)<sup>20</sup> | 55 | Male | USA | Costa Rica | 2015 | 8 | 3-12 | <39 | – |
| Duffy (2009)<sup>4</sup> | – | Female | USA | Yap Island | 2007 | 13 | 7-21 | <34 | – |
| Fonseca (2014)<sup>21</sup> | – | Female | Canada | Thailand | 2013 | 16 | 1-18 | <24<sup>d</sup> | 26-28 |
| Foy (2011)<sup>22</sup> | 36 | Male | USA | Senegal | 2008 | 24 | 5-30 | <33 | <33<sup>e</sup> |
|  | 27 | Male | USA | Senegal | 2008 | 24 | 4-29 | <33 | <33<sup>e</sup> |
|  | – | Female | USA | USA<sup>f</sup> | 2008 | 7 | 3-11 | 15-34 | <16<sup>e</sup> |
| Ginier (2016)<sup>23</sup> | 51 | Female | Switzerland | Guatemala, El Salvador | 2015 | 14 | 3-18 | <24 | >23 |
| Gyurech (2016)<sup>24</sup> | 44 | Female | Switzerland | Brazil | 2015 | 1 | 4-17 | 19-23 | <23 |
| Korhonen (2016)<sup>25</sup> | 37 | Male | Finland | Maldives | 2015 | 183 | 1-185 | – | <191<sup>g</sup> |
| Kutsuna (2014)<sup>26</sup> | Early 30s | Female | Japan | Bora Bora | 2013-2014 | 10 | 5-16 | <21 | <21<sup>h</sup> |
| Kwong (2013)<sup>27</sup> | 52 | Female | Australia | Indonesia | – | 9 | 0-10 | – | 13-24 |
| Leung (2015)<sup>28</sup> | 27 | Male | Australia | Indonesia<sup>i</sup> | – | 6 | 2-9 | – | <14<sup>j</sup> |
| Maria (2016)<sup>29</sup> | 60s | Female | France | Martinique Island | 2015 | 22 | 1-24 | <28 | – |
|  | 20s | Male | France | Brazil | 2015-2016 | 8 | 0-9 | <17 | – |
|  | 50s | Male | France | Colombia | 2015-2016 | 29 | 0-30 | 31-37 | –<sup>k</sup> |
| Shinohara (2016)<sup>30</sup> | Early 40s | Male | Japan | Thailand | 2014 | 7 | 1-9 | 10-14 | >10<sup>d</sup> |
| Simpson (1964)<sup>31</sup> | 28 | Male | Europe | Uganda | – | 76 | 0-77 | <78 | >2<sup>c</sup> |
| Summers (2015)<sup>32</sup> | 48 | Male | USA | Ecuador, Peru, Bolivia, Chile, Easter Island, French Polynesia, Hawaii | 2013 | 34 | 0-35 | <45 | – |
| Tappe (2015)<sup>33</sup> | 45 | Female | Germany | Malaysia | 2014 | 22 | 5-28 | 29-33 | <30 |
| Tappe (2014)<sup>34</sup> | Early 50s | Male | Germany | Thailand | 2013 | 12 | 0-12 | <22 | <22 |
| Wæhre (2014)<sup>35</sup> | 31 | Female | Norway | Tahiti | 2013 | 15 | 0-16 | 20-52 | 20-52<sup>l</sup> |
| Zammarchi (2015)<sup>36</sup> | Early 60s | Male | Italy | Brazil | 2015 | 12 | 0-13 | <16 | <16 |
| Zammarchi (2015)<sup>37</sup> | Early 30s | Female | Italy | French Polynesia | 2013-2014 | 19 | 0-20 | 22-58 | >22 |
|  | Early 30s | Male | Italy | French Polynesia | 2013-2014 | 19 | 0-20 | 22-56 | <23 |

<sup>a</sup> Days to viral clearance in sera
<sup>b</sup> Volunteer inoculation of Zika virus
<sup>c</sup> Viral shedding determined from mouse inoculation
<sup>d</sup> Equivocal result counted as positive
<sup>e</sup> Serum was positive by PCR and negative by culture
<sup>f</sup> Probable sexual transmission
<sup>g</sup> Serum was negative by PCR; urine was positive by PCR at later visit
<sup>h</sup> Serum was negative by PCR; urine was positive by PCR
<sup>i</sup> Possible transmission from monkey bite or mosquito
<sup>j</sup> Serum and swab of monkey bite site were negative by PCR; nasopharyngeal swab was positive by PCR
<sup>k</sup> No sera tested; plasma and urine positive by PCR; plasma was negative by PCR and urine and saliva were positive by PCR at a later visit
<sup>l</sup> Serum was positive by PCR and negative by culture

Of the cases in the final data set, 23 were infected while traveling in endemic areas, one via sexual transmission, and one through experimental infection. The vast majority of cases occurred after 2008 and were among residents of the United States or Europe (Table 1). None of the reported infections were among children, and there were roughly equal numbers of males and females (14/25 male).

### Key Distributions

We estimate the median incubation period of Zika virus to be 5.9 days (95% CI: 4.4-7.6), with a dispersion of 1.46 (95% CI: 1.23-1.94). Hence, 5% of cases will develop symptoms by 3.2 days after infection (95% CI: 1.7-4.6), 25% by 4.6 days (95% CI: 3.1, 6.0), 75% by 7.6 days (95% CI: 5.8-10.4), and 95% by 11.2 days (95% CI: 7.6-18.0) (Figure 2A).

**Figure 2:**
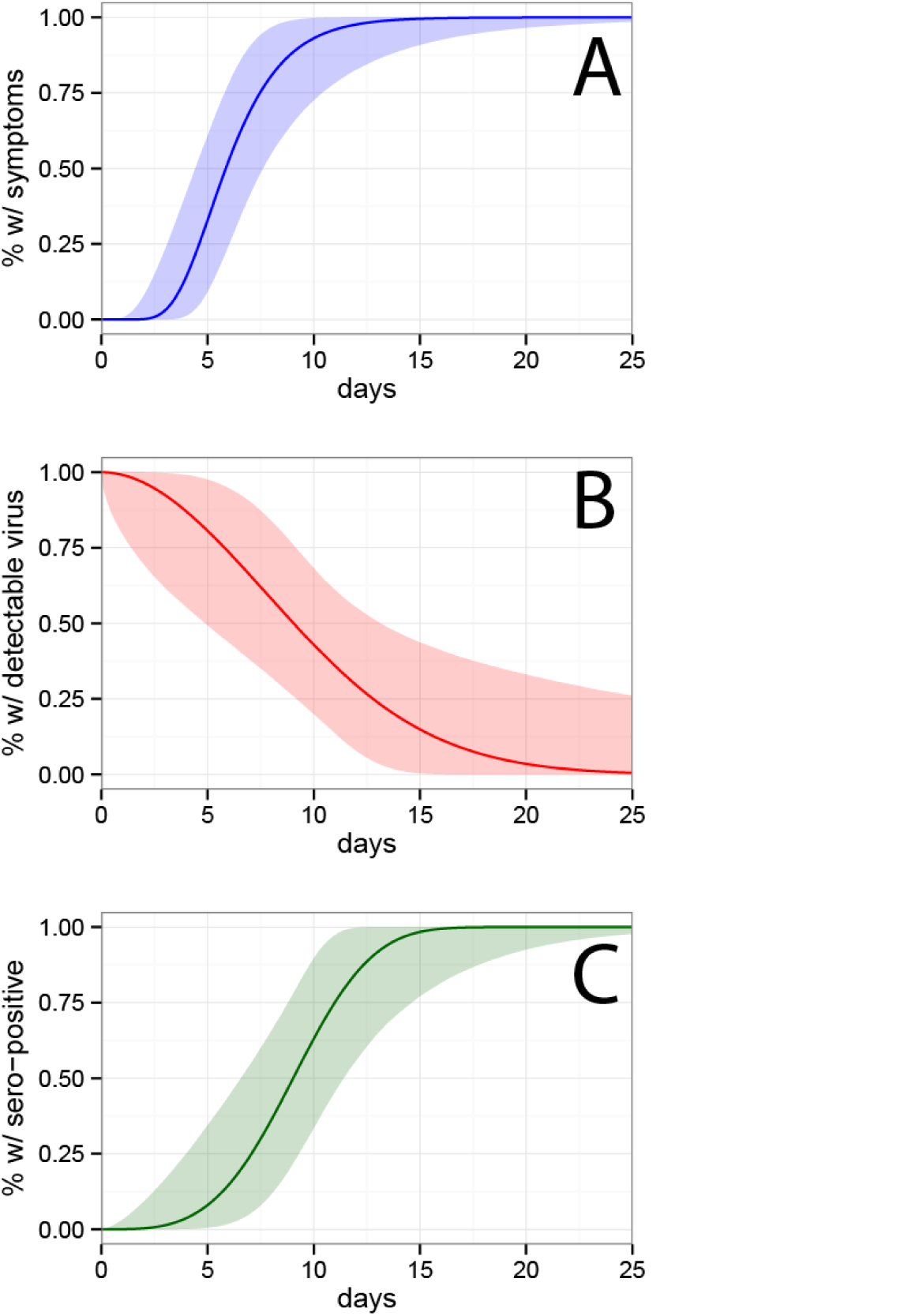
Percentage of the population with **(A)** symptom onset after a given time, **(B)** still shedding at a given day, and **(C)** testing seropositive as of a given day. Shaded regions indicate 95% credible intervals.

We estimate the mean time to viral clearance, defined as having no detectable virus in the blood, to be 9.9 days (95% CI: 6.9-21.4). We estimate that 5% of cases will have no detectable virus by 2.4 days after infection (95% CI: 0.09-5.9), 25% by 5.8 days (95% CI: 1.4, 9.2), 75% by 12.7 days (95% CI: 9.2-25.9), and 95% by 18.9 days (95% CI: 13.6-79.4) (Figure 2B).

We estimate the mean time to seroconversion is 9.1 days (95% CI: 7.0-11.6).). We estimate that 5% of cases will have detectable antibodies by 4.4 days after infection (95% CI: 1.3-7.0), 25% by 7.1 days (95% CI: 4.0, 9.2), 75% by 10.1 days (95% CI: 8.7-14.6), and 95% by 13.7 days (95% CI: 10.6-21.7) (Figure 2C).

### Implications for Surveillance and Blood Supply Safety

The mean time to viral clearance from the blood is 9.9 days, hence, in settings with ongoing transmission, if no screening of any type were performed, there would be a 9.9 per 100,000 donors increase (95% CI: 6.9-21.4) in the risk of a blood donation being infected with Zika for every 1 in 100,000 increase in daily Zika incidence. Preventing those with recent symptoms of possible Zika infection from donating would only decrease this risk by 7% (RR 0.93, 95% CI 0.89-0.99), as 80% of individuals with Zika infection are asymptomatic, and even those who do develop symptoms will be infectious but asymptomatic for an average of six days (assuming blood donations can transmit Zika virus from the moment of infection). Serological screening is more effective, reducing the risk by 29% (RR 0.71, 95% CI: 0.28-0.88), but still only marginally improves blood supply safety.

Since it may not be practical to stop blood donations until the Zika epidemic has passed, countries may consider virologic (i.e., nucleic acid) testing of particular lots of donated blood for targeted use in pregnant women. Still, even nucleic acid testing is imperfect; we did find a single case of a negative virologic blood test followed by a positive one, though this was in the context of a perinatal transmission and not part of our main analysis.^38^

In settings where the risk is solely from imported Zika cases, ensuring blood supply safety is far easier. By 23.4 (95% CI, 14.3-154.3) days after infection, 99% of infections are expected to no longer have detectable virus in their blood. While this number cannot be estimated with confidence given the low number of observations it is based upon, it can serve as the basis for a risk averse donation rule (e.g. no donation for 300 days after travel to a Zika endemic regions, over twice the upper limit of the confidence interval for this estimate).

It is important to note that here we assume that not having detectable virus in blood implies safe blood donation; however, risk to the blood supply when virus is present in other fluids cannot be ruled out. We found four cases in which virus was no longer detectable in blood but a saliva, nasal, or urine sample tested positive (Table 1). While we have inadequate data to estimate the time to viral clearance in these fluids, we estimate the latest of these positive tests was 12.0 days after infection (95% CI: 10.1-18.2) for the individuals in our dataset. Duration of viremia in other fluids may be relevant to other public health recommendations (e.g., how long to abstain from sex with a potentially pregnant partner).

## DISCUSSION

As of time of writing, the WHO reports the incubation period of Zika virus as unclear, but likely “a few days.”^39^ Likewise, the US Centers for Disease Control and Prevention (CDC) states that the incubation period of Zika is unknown but probably “a few days to a week,”^40^ and the European Centre for Disease Prevention and Control (ECDC) estimates 3-12 days.^41^ Our analysis substantially clarifies the true incubation period for Zika virus infection and the amount of uncertainty that remains. We similarly illuminate the distribution of time to seroconversion and time to viral clearance.

Understanding what is known about key distributions in the natural history of Zika virus infection is an important component of designing and evaluating screening and surveillance protocols, as we illustrate with an analysis of screening for Zika infection in blood donors. While the risk is quite low, it scales with Zika incidence, which in turn is hard to measure due to the high number of asymptomatic cases. Screening is important, but only a direct antigen test can have any hope of substantially reducing risk, though serologic tests may be able to offer a marginal (~30%) improvement.

This analysis is based on published data that was collected for reasons other than estimation of these key distributions; as such we were required to make several assumptions. We assumed that the virologic testing of blood or sera is 100% sensitive for detecting Zika virus; however there is evidence that viral shedding can continue far longer in urine and other bodily fluids, raising concerns that virus may exist in the blood below the limit of detection. We assumed that the distribution of time to seropositivity is independent of previous infection with other flaviviruses (those with prior flavivirus infections will likely seroconvert more quickly). Since the majority of the cases included in our analysis were travelers returning to countries with little endemic flavivirus circulation, it is likely our estimates of time to seroconversion are conservative (i.e., long). Further, the majority of our data comes from presumed mosquito infections, and these distributions may differ for other routes of infection (e.g., perinatal, sexual). Likewise, all of the cases we report were symptomatic, and the distribution of time to seroconversion and viral clearance may differ in asymptomatic individuals. However, the biggest limitation of our analysis is the small number of cases, which both increases uncertainty and the potential for bias.

Despite the limitations of this analysis, our estimates are the most detailed, quantitative estimates to date for the natural history of Zika virus. These estimates can be used to target surveillance in both endemic settings and for returning travelers as well as guide empirical efforts to study basic features of this pathogen.

## ACKNOWLEDGEMENTS

We would like to thank Prof. Alfonso Javier Rodriguez-Morales for providing additional data on reported cases. We also thank Joshua Sharfstein for helping to bring together the team to perform this rapid review. We further thank the Department of Epidemiology and the Office of Public Health Practice and Training at the Johns Hopkins Bloomberg School of Public Health for financially supporting this work.

